# Dopamine desynchronizes the retinal clock through a melanopsin-dependent regulation of acetylcholine retinal waves during development

**DOI:** 10.1101/2021.04.23.441100

**Authors:** Chaimaa Kinane, Hugo Calligaro, Antonin Jandot, Christine Coutanson, Nasser Haddjeri, Mohamed Bennis, Ouria Dkhissi-Benyahya

## Abstract

Dopamine (DA) plays a critical role in retinal physiology, including resetting of the retinal circadian clock that in turn regulates DA release. DA acts on major classes of retinal cells by reconfiguring electrical and chemical synapses. Although a bidirectional regulation between intrinsically photosensitive melanopsin ganglion cells (ipRGCs) and dopaminergic cells has been demonstrated during development and adulthood, DA involvement in the ontogeny of the retinal clock is still unknown.

Using wild-type *Per2*^*Luc*^ and melanopsin knockout (*Opn4*^*-/-*^::*Per2*^*Luc*^) mice at different postnatal stages, we found that the retina can generate self-sustained circadian rhythms from postnatal day 5 that emerge in the absence of external time cues in both genotypes. Intriguingly, DA lengthens the endogenous period only in wild-type retinas, suggesting that this desynchronizing effect requires melanopsin. Furthermore, blockade of cholinergic retinal waves in wild-type retinas induces a shortening of the period, similarly to *Opn4*^*-/-*^::*Per2*^*Luc*^ explants. Altogether, these data suggest that DA desynchronizes the retinal clock through a melanopsin-dependent regulation of acetylcholine retinal waves, thus offering a new role of melanopsin in setting the period of the retinal clock during development.

## Introduction

Dopamine (DA) is a key neurotransmitter, synthesized and released by a sparse population of retinal amacrine cells. Hence, DA is implicated in daily and circadian retinal functions, including phase resetting of the retinal circadian oscillator (Ruan et al., 2008), adaptation to light, clock gene regulation (Jackson et al., 2012; Witkovsky, 2004; Yujnovsky et al., 2006), photoreceptor, amacrine and ganglion cell coupling (Arroyo et al., 2016; Hampson et al., 1992; Ribelayga et al., 2008) and eye development (Zhou et al., 2017).

Besides, DA triggers a wide array of intracellular changes via two classes of dopaminergic receptors: the D1-type (D1R and D5R) are coupled to Gα_s/olf_ and stimulate the production of cAMP and the activity of protein kinase A, whereas the D2-type (D2R, D3R, and D4R) are coupled to Gα_i/o_ and negatively regulate cAMP, resulting in a decrease in protein kinase A activity (Seeman and Van Tol, 1994). Most retinal cells, including Müller glial cells, can be modulated by DA as they express at least one type of dopaminergic receptors. In rodents, rods and cones express D4Rs and may express D2Rs in some species (Goyal et al., 2020; Witkovsky, 2004), whereas horizontal, bipolar, amacrine, and ganglion cells mainly express D1Rs (Derouiche, 1999; Farshi et al., 2016; Nguyen-Legros et al., 1997; Popova, 2014; Travis et al., 2018; Veruki, 1997). In the adult, DA is assumed to act on the retinal circadian clock through D1Rs, whereas D2Rs play a role in the induction of the *Per* clock gene by light (Jackson et al., 2012; Ribelayga and Mangel, 2003; Ribelayga et al., 2008; Yujnovsky et al., 2006).

Despite the description of retinal dopaminergic neurons several decades ago, their roles in the development of the retinal circadian clock functioning are poorly understood. The mammalian retinal clock is composed of an integrated network of cellular clocks localized throughout the retina (Dkhissi-Benyahya et al., 2013; Dorenbos et al., 2007; Jaeger et al., 2015; Ruan et al., 2008; Schneider et al., 2010) that give rise to the rhythmic regulation of retinal physiology and function in the adult. Even if clock gene expression has been recently reported at embryonic stages in the mouse retina (Bagchi et al., 2020), remarkably, it is still unknown when retinal cells first express circadian rhythms and how they synchronize during development. DA synthesis and release are regulated by the retinal circadian clock (Buonfiglio et al., 2014; Doyle et al., 2002; Felder-Schmittbuhl et al., 2018, 2017; Iuvone et al., 1978). Moreover, dopaminergic cell activity has been shown to be stimulated by rods, cones, and/or melanopsin intrinsically photosensitive retinal ganglion cells (ipRGCs) upon illumination (Pérez-Fernández et al., 2019; Zhang et al., 2007; Zhao et al., 2017). In turn, ipRGCs stratify with dopaminergic cells during development and adulthood (Renna et al., 2015; Vugler et al., 2007) to convey retrograde signalling (Prigge et al., 2016; Wang et al., 2008). Furthermore, Munteanu and colleagues suggested that melanopsin does not play a critical role in setting dopaminergic cell number and DA levels in the developing mouse retina (Munteanu et al., 2018). However, we have previously shown that the adult melanopsin knockout mouse (*Opn4*^*-/-*^) exhibits a dysfunction of the retinal dopaminergic system and a shortening of the endogenous period of the retinal clock (Calligaro et al., 2019; Dkhissi-Benyahya et al., 2013). Interestingly, during development, a bidirectional regulation between ipRGCs and cholinergic retinal waves has been suggested (Arroyo et al., 2016; Kirkby and Feller, 2013). Particularly, ipRGCs were shown to modulate the circuitry of cholinergic retinal waves in the neonatal retina even in darkness (Renna et al., 2011). These waves consist of spontaneous periodic bursts of ganglion cell activity (Ackman et al., 2012; Wong et al., 1993), observed from postnatal day 1 (P1) to P9- P11 (Blankenship and Feller, 2010; Feller et al., 1996). In turn, in the absence of cholinergic waves, dopaminergic signalling modulates the extent of ipRGCs gap junction coupling in the developing retina (Arroyo et al., 2016; Kirkby and Feller, 2013).

In the current study, we first established the onset and the fundamental features of the retinal clock in the *Per2*^*Luc*^ wild-type and the *Opn4*^*-/-*^::*Per2*^*Luc*^ mice during development and then determined whether DA plays a crucial role in retinal clock ontogeny. We then analyzed whether the absence of melanopsin would affect the maturation of the retinal clock during development.

## Results

### First detection of retinal PER2::Luc circadian oscillations *in vitro* at postnatal day 5

Retinal explants from embryonic day 18 (E18), different postnatal stages (P1, P5, P8, P11, P15, P30), and from adult (P60-90) *Per2*^*Luc*^ mouse were continuously recorded for at least six days without medium change (Figure 1A-2A; Supplementary Figure 1). A first rhythmic expression of PER2::Luc was observed at P5 after 2 days *in vitro*, which became more robust, with higher amplitude from P8. During retinal maturation, the endogenous period was significantly shortened between P8 and P11 (respectively 26.35 ± 0.12 h and 25.54 ± 0.07 h; p < 0.001), and between P15 and P30 (respectively 25.71 ± 0.24 h and 24.55 ± 0.08 h; p < 0.001; Figure 1B). Then, it was lengthened between P30 and adulthood (25.19 ± 0.14 h; p ≤ 0.01). The phase of PER2::Luc expression was constant from P5 (CT 11.58 ± 1.83) to P11 (CT 13.05 ± 0.35; p ≥ 0.05), then was progressively delayed between P11 and P15 (CT 16.54 ± 0.39; p ≤ 0.001), and from P15 to P30 (CT 20.40 ± 0.65; p ≤ 0.001). No significant difference was detected between P30 and the adult (CT19.57 ± 0.61; p ≥ 0.05). The amplitude of PER2::Luc oscillations increased from P5 to P30 (P5:12.27 ± 1.14 cps; P8: 30.51 ± 2.04 cps; P11: 56.34 ± 4.58 cps; P15: 267.60 ± 83.42 cps; and P30: 604.80 ± 68.50 cps), followed by a significant reduction between P30 and the adult (143.43 ± 23.85 cps; p ≤ 0.001).

**Figure 1:**
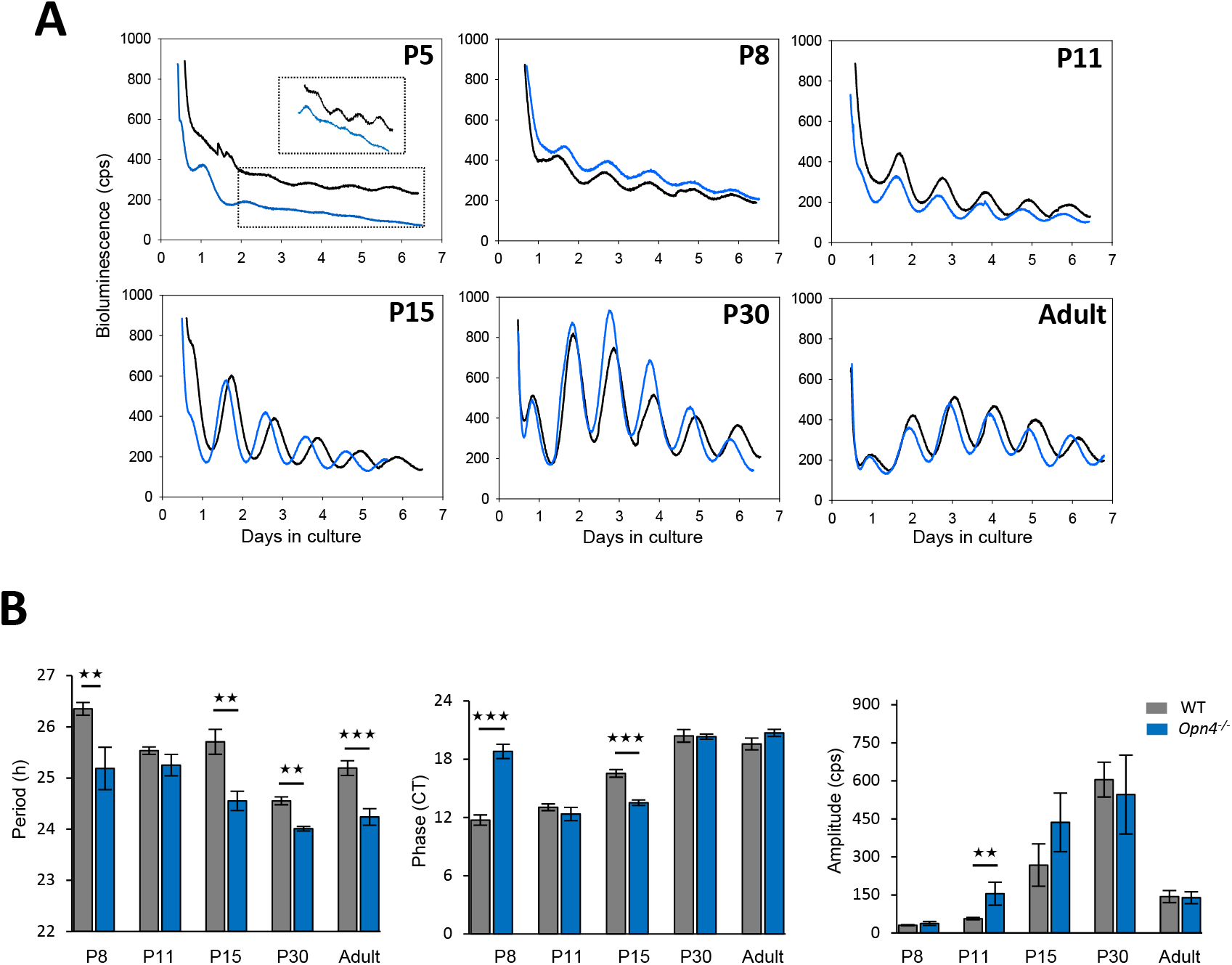
Ontogenesis of PER2::Luc circadian oscillations in wild-type and *Opn4*^*-/-*^ retinal explants. **A**- Representative bioluminescence recording of PER2::Luc retinal explants from *Per2*^*Luc*^ (black line) and *Opn4*^*-/-*^*:: Per2*^*Luc*^ (blue line) mice at different postnatal stages (P5, P8, P11, P15, P30) and in the adult. Retinal explants were cultured for at least 6 days without medium change. The dotted black rectangle corresponds to an enlargement of bioluminescence traces of both genotypes at P5 **B-** Means of the endogenous period, the circadian phase, and the amplitude in both genotypes during development. The total numbers of retinas used were respectively for wild-type mice: P8, n=10; P11, n=10; P15, n=6; P30, n=8; adult, n=10 and for *Opn4*^*-/-*^*::Per2*^*Luc*^; P8, n=6; P11, n=6; P15, n=7; P30, n=6 and adult, n=10. Bars represent the mean ± SEM (ANOVA, ** = p <0.01; *** = p <0.001).

### Impact of the absence of melanopsin on the ontogeny of the retinal clock

We previously described a loss of clock gene rhythms in the photoreceptor layer of the *Opn4*^*-/-*^ mouse (Dkhissi-Benyahya et al., 2013). Hence, we decided to analyze the impact of the absence of melanopsin on the retinal clock’s fundamental features. Similar to the wild-type mouse, PER2::Luc oscillations started at P5 in the *Opn4*^*-/-*^::*Per2*^*Luc*^ mouse and became more robust with a higher amplitude from P8 (Figure 1A). The main difference between both genotypes concerned the period that was significantly shortened at all developmental stages (excepting P11) in *Opn4*^*-/-*^::*Per2*^*Luc*^ retinal explants (P8: 25.19 ± 0,41 h; P15:24.55 ± 0.19 h; adult: 24.24 ± 0.16 h; p≤0.01, Figure 1B). A significant delay and advance of the phase of PER2::Luc oscillations were respectively observed at P8 (CT 18.80 ± 0.73; p≤0.001) and P15 (CT 13.51 ± 0.28; p≤0.001) in retinal explants from *Opn4*^*-/-*^::*Per2*^*Luc*^ mouse compared to the wild-type mouse that was not observed in the adult. The amplitude of retinal PER2::Luc oscillations is mainly similar between genotypes at the different developmental stages.

### Spontaneous PER2::Luc oscillations develop *in vitro*

As circadian oscillations were undetectable before P5, and in order to determine whether rhythms could eventually develop *in vitro*, we cultured retinal explants from P1 *Per2*^*Luc*^ mice for 9 days without medium change (Figure 2A). These explants did not express rhythms for the first days in culture, then spontaneous oscillations with low amplitude were observed after 5 days *in vitro* (5-DIV P1) in all retinal explants. We thus compared the circadian parameters of 5-DIV P1 retinal explants with P5 cultured retinal explants (Figure 2B). No differences were observed in both the endogenous period (5-DIV P1: 27.22 ± 0.18 h, n=5; P5: 26.35 ± 0.62 h, n=6; p = 0.25), and the phase (5-DIV P1: CT 10.85 ± 1.36; P5: CT 11.58 ± 1.83; *p* = 0.76) between both conditions, whereas the amplitude was higher in P5 cultured explants (5-DIV P1: 4.92 ± 1.78 cps; P5: 12.27 ± 1.14 cps; p = 0.006).

**Figure 2:**
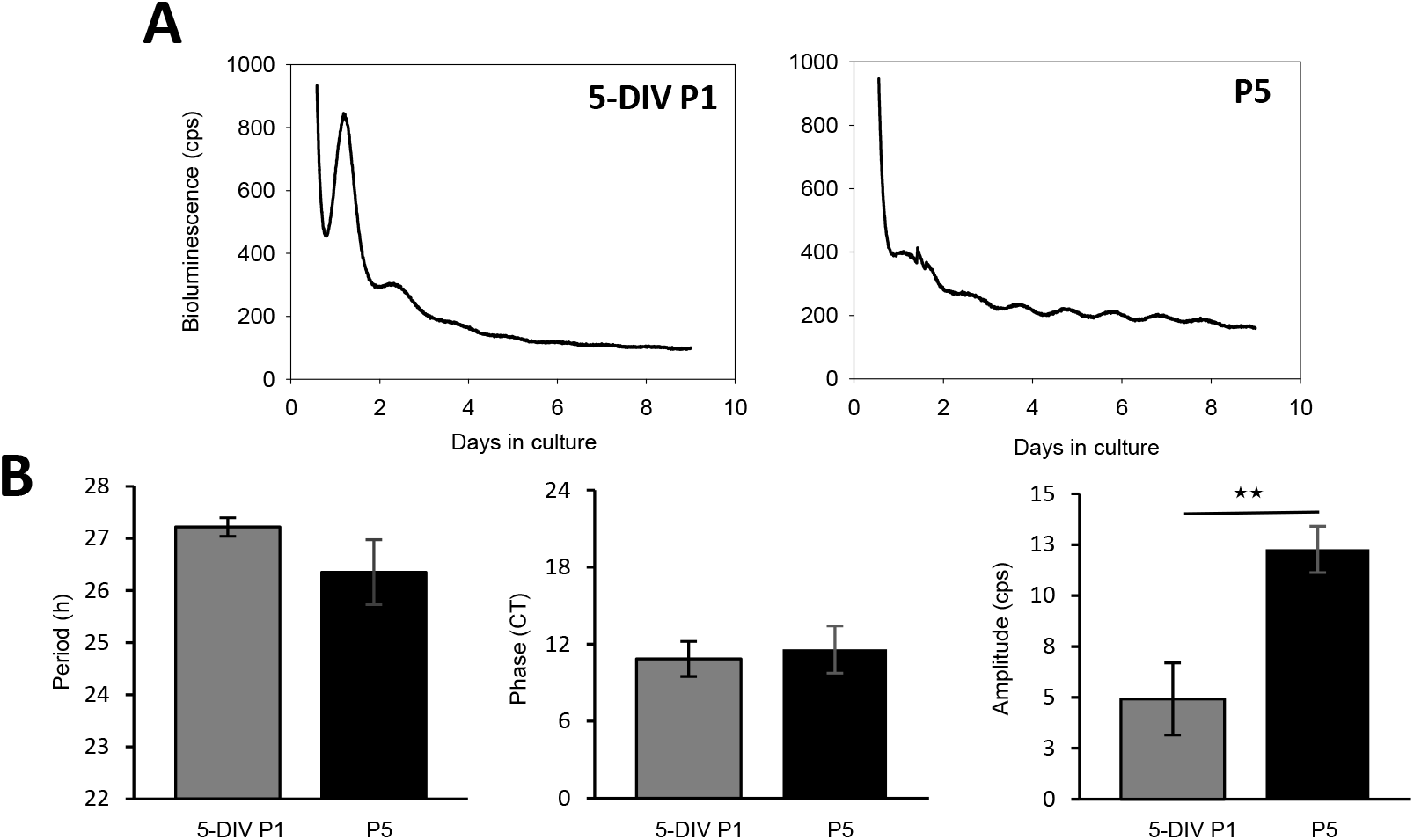
Spontaneous PER2::Luc oscillations develop *in vitro* from P1 mouse retinal explants. **A-** Representative bioluminescence recording of PER2::Luc from P1 and P5 retinal explants. After 5 days *in vitro*, P1 retinal explants (5-DIV P1) exhibit analysable PER2::Luc oscillations. **B-** Means of the endogenous period, the phase, and the amplitude of 5-DIV P1 and P5 PER2::Luc retinal explants. The analysis was performed on the 3 first complete oscillations for each explant. The total numbers of retinas used were: 5-DIV P1, n= 5, and P5, n=6. Bars represent the mean ± SEM (ANOVA, ** = p <0.01).

### Effect of DA on PER2::Luc retinal rhythms during development in the wild-type mouse

Despite the important role of DA in resetting the retinal circadian clock, its role in the ontogeny of this oscillator remains unknown. We first examined whether DA is required for the generation of PER2::Luc rhythms. Retinal explants were treated at P8, P15, and P30 with both L-AMPT and Res, an inhibitor of DA synthesis and of vesicular DA uptake, respectively, and compared to untreated retinas (Figure 3A). At the 3 developmental stages, PER2::Luc rhythms persisted after DA depletion, with no changes in the endogenous period at P8 and P15 (respectively 26.45 ± 0.23 h and 25.50 ± 0.25 h; (p ≥ 0.05) whereas a slight lengthening was observed at P30 (24.94 ± 0.11 h; p≤0.05) by comparison to non-depleted retinas of the same stage (p ≥ 0.05). DA depletion had also no effect on the phase of PER2::Luc oscillations at P8 (CT 11.51 ± 0.87), induced a small advance at P15 (DA-depleted retinas: CT 14.73 ± 0.27; controls: CT 16.54 ± 0.39; p≤0.05) and P30 (DA-depleted retinas: CT18.17 ± 0.50; controls: CT 20.40 ± 0.65; p≤0.05) whereas the amplitude was increased at P8 (74.72 ± 32.05 cps) and P15 (580.14 ± 30.50 cps). These results suggest that DA is not required for the generation of PER2::Luc rhythm during the mammalian retinal clock development. We then treated retinal explants from the same developmental stages (P8, P15, and P30) with DA, or Apo, a non-selective DA agonist (Figure 3A). An important result was that the supplementation in DA or Apo in P8 retinal explants significantly lengthened the endogenous period (DA: 27.17 ± 0.13 h; Apo: 27.54 ± 0.15 h, p≤0.01), delayed the phase (DA: CT 15.69 ± 1.01; Apo: CT 16.89 ± 0.50, p≤0.01) and increased the amplitude (DA: 177.98 ± 33.99 cps; Apo: 177.16 ± 25.28 cps) of PER2::Luc oscillations by comparison to non-supplemented explants. Both drugs also lengthened the period at P30 (DA: 25.53 ± 0.22 h; Apo: 26.76 ± 0.73 h, p≤0.01). The effect of DA supplementation on the amplitude of the retinal rhythm was the more variable, with an increase at P8 and P15 (p≤0.05) and a decrease at P30 (p≤0.01). We did not observe any effect of DA supplementation in the period of PER2::Luc oscillation (25.67 ± 0.43 h) in the adult (data not shown). We then investigated whether DA supplementation may affect the onset of spontaneous oscillations of PER2::Luc *in vitro*. As shown in supplementary Figure 2, P1 retinal explants could not express circadian oscillations in the presence of DA for the first 5 days in culture (5-DIV P1 + DA). Furthermore, as observed in the non-supplemented 5-DIV P1 retinas, the oscillations began to emerge after 5 days *in vitro* with very low amplitudes. No significant differences were observed in the period and the phase between 5-DIV P1 explants supplemented in DA and P5 retinal explants.

**Figure 3:**
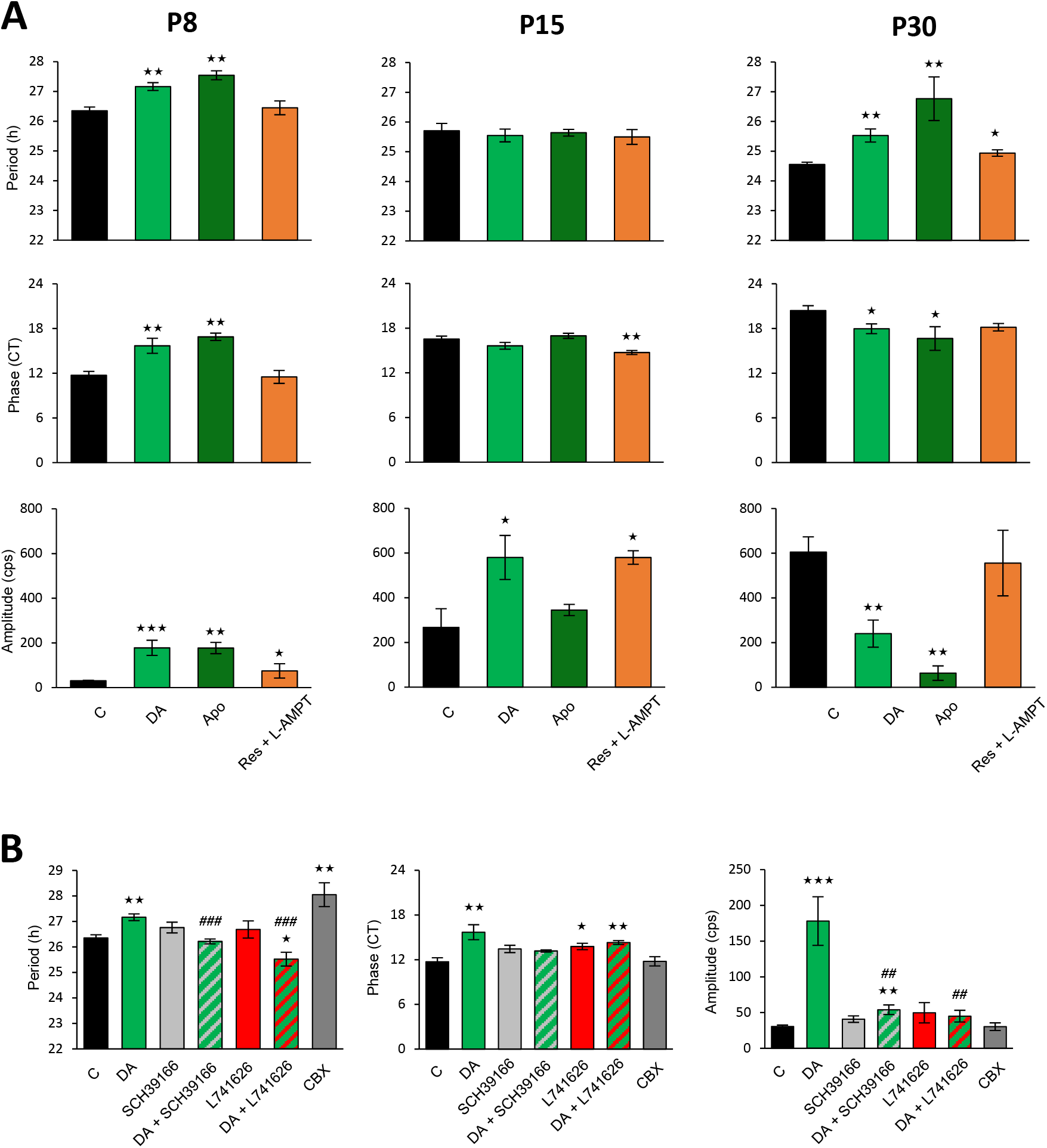
Dopamine desynchronizes the circadian retinal clock through D1 and D2 dopaminergic receptors during development. **A-**Means of the endogenous period, the phase and the amplitude of the retinal clock at 3 postnatal developmental stages (P8, P15 and P30). Retinal explants were supplemented with DA, Apo, or a combination of Res + L-AMPT and are compared to non- supplemented control retinas. The total numbers of retinas used were: P8: C, n=10; DA, n=7; Apo, n= 4; Res + L-AMPT, n= 8); P15: C, n=8. DA, n=6; Apo, n=3; Res + L-AMPT, n=5; P30: C, n=8; DA, n=9; Apo, n=5; Res + L-AMPT, n=6. **B-** Means of the endogenous period, the phase and the amplitude of P8 retinal explants, supplemented with DA (n=7), a D1R DA antagonist (SCH39166, n=6), a D2R DA antagonist (L741626, n=6), DA combined to SCH39166 (n=6) or L741626 (n=6), or a general gap junction blocker (CBX, n=4) and are compared to non-supplemented control retinas (C, n=10). Bars represent mean ± SEM (* = p <0.05; ** = p <0.01; *** = p <0.001, comparison with C group; ^##^ = p ≤ 0,01; ^###^ = p ≤ 0,001, comparison with DA group).

### The effect of DA on the retinal circadian clock is mediated by both D1 and D2 dopaminergic receptors and gap junctions

During development, we observed that the endogenous period of PER2::Luc oscillations was progressively shortened from P8 to P30 and that the application of DA lengthened this clock parameter. To determine which dopaminergic receptors mediate this effect, we focused on the P8 stage. Retinal explants were first treated with the D1R SCH39166 or the D2R L741626 antagonists to characterize their respective effects on the clock. As shown in figure 3B, the application of SCH39166 or L741626 at these concentrations did not change the period (respectively: 26.76 ± 0.21 h and 26.69 ± 0.34 h) and the amplitude, with a slight delay of the phase only with L741626 (CT13.77 ± 0.43, p = 0.04) compared to control retinas. We further combined DA treatment with SCH39166 or L741626. Both combinations blocked or reduced the effect of DA on the period (p≤0.01) of PER2::Luc oscillations. In addition, the period lengthening was also observed when P8 retinal explants were treated with carbenoxolone (CBX, 100 µM), a general gap junction blocker (Figure 3B; p≤0.01).

### DA does not alter the period of PER2::Luc retinal rhythms in the *Opn*_*4*_^*-/-*^ mouse during development

Since the supplementation of DA at P8 and P30 lengthened the period of PER2::Luc rhythms in wild-type retinal explants, we analyzed the effect of DA in retinal explants from *Opn4*^*-/-*^::*Per2*^*Luc*^ mouse at both stages. Contrary to what had been observed in the wild-type retinal explants, in the melanopsin knockout explants, DA did not lengthen the period and advanced the phase at P8 (Figure 4; CT 14.50 ± 0.28; p≤0.01). In both genotypes, the amplitude was increased at P8 (p≤0.01). Then, we assessed whether the absence of DA effect on the period could be related to the differential relative expression of D1- (D1 and D5 receptors)- and D2- like (D2 and D4) receptors at P8 and P30 (Figure 5). At P8, no difference was observed in the relative expression of both D1- and D2-like receptors between genotypes, whereas at P30, expressions of D1R and D2R mRNAs were significantly increased in the retinas of *Opn4*^*-/-*^ mouse compared to wild-type mouse (for D1R; wild-type: 1.15 ± 0.05 and *Opn4*^*-/-*^: 1.65 ± 0.06, p <0.01; for D2R; wild-type: 2.18 ± 0.17 and *Opn4*^*-/-*^: 3.90 ± 0.23, p <0.01). No differences were observed for D4Rs and D5Rs at both stages. Finally, if cholinergic retinal waves are involved in the setting of the retinal period through DA, blocking them in wild-type retinal explants should result in a pattern similar to what we observed in *Opn4*^*-/-*^ explants. As predicted, the blockade of retinal waves by MMA, an antagonist of nicotinergic receptors, significantly shortened the period and delayed the phase of PER2::Luc oscillations in P8 wild-type retinal explants (Figure 6).

**Figure 4:**
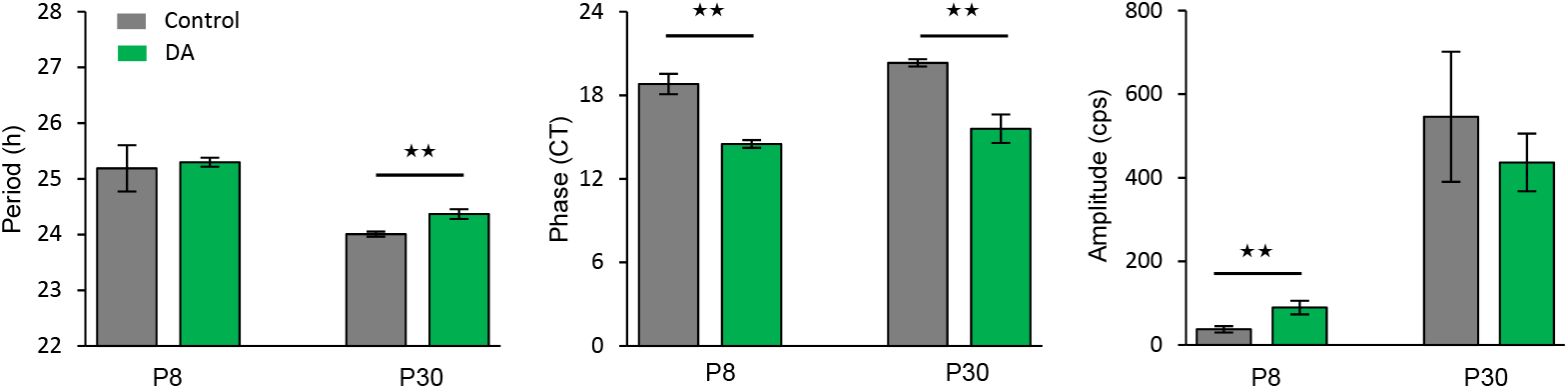
Effect of dopamine on the retinal clock in the absence of melanopsin. Means of the endogenous period, the phase, and the amplitude of P8 and P30 retinal explants from *Opn4*^*-/-*^*::Per2*^*Luc*^ mice, supplemented with DA and compared to non- supplemented control retinas (C). The total numbers of retinas used were: P8: C, n=6; DA, n=4; P30: C, n=5; DA, n=5. Bars represent mean ± SEM (ANOVA, ** = p <0.01).

**Figure 5:**
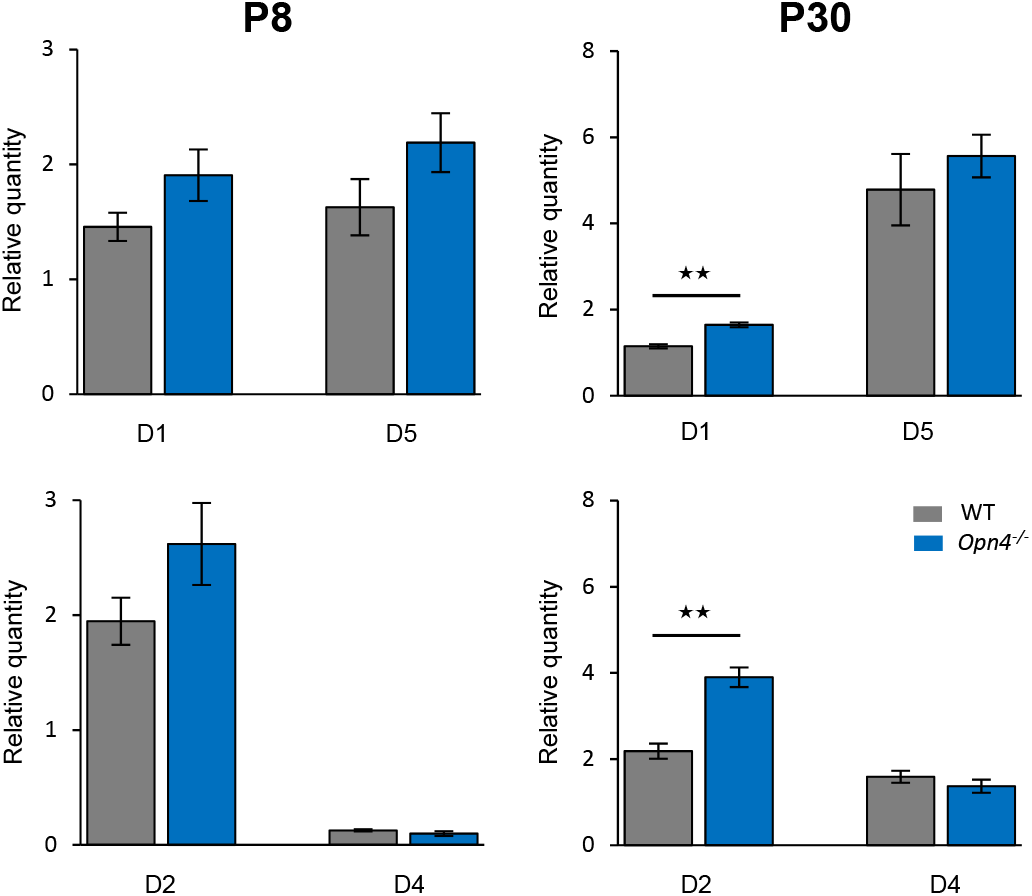
Relative expressions of D1- and D2-like dopaminergic receptors are similar between wild-type and *Opn4*^*-/*-^ retinas at P8. Means of the relative expression of D1-like (D1, D5) and D2-like (D2, D4) receptors in the retinas of wild-type (WT, grey bars) and *Opn4*^*-/-*^ (blue bars) mice at 2 developmental stages, P8 and P30. The total numbers of retinas used were: WT, n=6 for P8 and P30; *Opn4*^*-/-*^, n=7 for P8 and n=4 for P30. Bars represent mean± SEM (ANOVA, ** = p <0.01).

**Figure 6:**
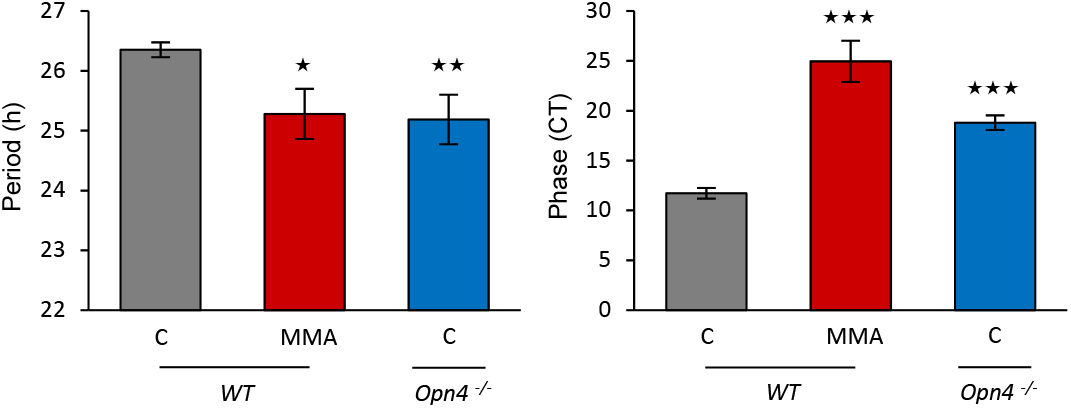
The blockade of acetylcholinergic waves shortens the endogenous period of the retinal clock in P8 explants of *Per2*^*luc*^ mice. Means of the endogenous period and the phase in control (C) wild-type (WT) and *Opn4*^*-/-*^::*Per2*^*luc*^ retinas and in WT retinas supplemented with mecamylamine acid (MMA). The total numbers of retinas used were: WT: C, n=10: MMA, n=6; *Opn4*^*-/-*^: C, n=8. Data from WT and *Opn4*^*-/-*^*::Per2*^*Luc*^ control retinas are already presented in figure 1B. Bars represent mean ± SEM (ANOVA, * = p <0.05; ** = p <0.01; *** = p <0.001).

## Discussion

In the mouse SCN, *in vivo* detectable fetal rhythms have been reported for *Per1* mRNAs at E17 and E18 for PER1 and PER2 proteins (Ansari et al., 2009; Shimomura et al., 2001), whereas *in vitro* studies detected PER2::LUC oscillations earlier at E13 (Landgraf et al., 2015) or E15 (Wreschnig et al., 2014). In the present study, we detected the first weak PER2::Luc retinal oscillations at P5, which can be considered a critical time in the maturation of the retinal clock. *Per2* expression was previously described in the inner part of the neuroblastic rat retina at P1 (Matějů et al., 2010), and a recent transcriptomic study reported that *Per2* is already expressed at E13 with a peak at P0 in the mouse retina (Bagchi et al., 2020). Thus, we cannot exclude that before P5, distinct cell populations may express PER2::Luc rhythms with different phases (Dkhissi-Benyahya et al., 2013), resulting in low amplitude rhythmicity difficult to detect at the level of the whole retina. From which retinal cells do circadian oscillations emerge? During development, the rhythmic bioluminescence signal could arise from different cell populations. In mice, retinal neurogenesis occurs between E11 and P10, prior to eye-opening at P14. Early progenitor cells that exit mitosis first generate ganglion and horizontal cells, along with cone photoreceptors and some amacrines while later progenitor cells give rise to late-born amacrine, rod, bipolar, and Müller glial cells (Bassett and Wallace, 2012; Belliveau and Cepko, 1999; Young, 1985). At P5, ganglion cells and cones are already present, and the inner part of the neuroblastic retina will give rise to the inner nuclear layer (INL) that becomes functional at the end of the first postnatal week (Sernagor et al., 2001). Thus, we can assume that the signal first arises from the inner retina since the INL was reported as the primary source of rhythmic PER2::Luc signals in the adult retina (Ruan et al., 2008).

Another important question is when the retina is able to sustain rhythms without maternal influence. This issue has been addressed by examining retinal rhythms over time in culture. When retinal explants obtained from P1 *Per2*^*Luc*^ mice are left in culture, they began to express spontaneous circadian oscillations *in vitro* around 4-5 days after culture’s beginning. This suggests an autonomous developmental program that ensues in isolated retinas without the influence of external signals. Accordingly, we found that the endogenous period and the phase of 5-DIV P1 and P5 retinal explants were similar, suggesting that the timing of the development of PER2::Luc oscillations is roughly identical between *in vivo* and *in vitro* at least for the first 5 days period in culture. For the SCN, contradictory results have been observed. One study reported that even if SCN explants did not initially show PER2::Luc rhythms at E14, then they express them after 4-5 days *in vitro* (Wreschnig et al., 2014), whereas others did not find evidence that SCN becomes circadian spontaneously *in vitro* (Carmona-Alcocer et al., 2018; Landgraf et al., 2015).

The retinal clock then develops gradually, with, in particular, a shortening of the endogenous period in both wild-type and *Opn4*^*-/-*^ mouse suggesting that cellular/layer oscillators progressively amplify and synchronize their genetic oscillations to drive coherent rhythms at the level of the whole retina. Indeed, cultures of isolated retinal layers or dissociated cells exhibit a lengthening of the period by comparison to whole retinal explants in the adult (Jaeger et al., 2015). Circadian oscillations are expressed at P5, an age when few synapses are present in the mouse retina. Indeed, first synapses in the IPL and the OPL appear in postnatal ages and continues for several weeks, reaching their peak at P21 (Tian, 2004), suggesting that other mechanisms could mediate synchrony before synapse formation. With a blocker of gap junctions, our results indicate that they play a role in establishing the onset of synchrony during the early stages of retinal clock development by unknown mechanisms. Besides, we have recently reported that the invalidation of melanopsin in the adult mouse shortens the endogenous period of the retinal clock (Calligaro et al., 2019), an alteration that was already observed when the first PER2::Luc oscillations are detected in the *Opn4*^*-/-*^ mouse. This implicates that melanopsin is involved in regulating the endogenous period of the retinal clock with a still non elucidated mechanism.

A good candidate for mediating the coupling between retinal cells during development is DA. Indeed, this neurotransmitter is released by amacrine cells whose processes make contacts within the inner and outer plexiform layers and has been shown to activate the transcriptional activity of the clock (Yujnovsky et al., 2006). In rodent retina, dopaminergic amacrine cells develop from the first days after birth, with a low DA level before P6 that rapidly increases in the following days (Martin-Martinelli et al., 1989; Wulle and Schnitzer, 1989). Surprisingly, while coupling within the adult mouse retinal clock did not rely on DA, the supplementation of DA or CBX early during the development significantly lengthen the period of PER2::Luc oscillations in the wild-type mice. This DA effect was blocked either by D1R- or D2R-like antagonists. The involvement of both DA receptor families was not surprising since DA was well known to modulate the phosphorylation of gap junctions by activating different intracellular pathways leading to an increase or decrease of their conductance across diverse neurons in the adult retina (Bloomfield and Volgyi, 2009). Thus, our result suggests that DA modulates the fundamental features of the retinal clock by changing the extent of gap junction coupling in the developing retina as recently suggested (Arroyo et al., 2016; Caval-Holme and Feller, 2019). The most striking result concerns the fact that DA did not lengthen the period of *Opn4*^*-/-*^ retinal explants but only at P8, suggesting that melanopsin expression is necessary for the desynchronizing effect of DA on the retinal clock at this stage. At P8, ipRGCs are the only functional photoreceptors, and even in the absence of light, these cells have been shown to modulate the plasticity of retinal cholinergic wave circuitry in the neonatal retina (Renna et al., 2011) via DA release (Arroyo et al., 2016; Kirkby et al., 2013). In turn, retinal waves regulate the network of developing ipRGCs that are mainly gap junction coupled to different neurons, including other ipRGCs (Arroyo et al., 2016). In the current study, the blockade of cholinergic retinal waves during the first postnatal week in wild-type retinal explants induced a pattern similar to the one observed in the *Opn4*^*-/-*^ with a shortening of the period and a delayed phase of the retinal clock. Accordingly, Arroyo and colleagues (2016) showed that this blockade dramatically increased the extent of ipRGCs coupling. Thus, the following model can be proposed (Figure 7). In wild-type C57Bl6 mouse, retinal DA is not circadian, and DA release is strictly light-driven (Iuvone et al., 1978). DA supplementation functions as a chemical signal for light, which in turn decreases the extent of gap junction coupling (Bloomfield and Volgyi, 2009; O’Brien and Bloomfield, 2018) between ipRGCs and other retinal cells, leading to a lengthening of the period of the retinal clock. Upon cholinergic blockage in the wild-type mouse, DA release is decreased, causing a robust electrical coupling between cells and a shortening of the period. The contribution of ipRGCs to retinal wave generation and dynamic is altered in the *Opn4*^*-/-*^ (Chew et al., 2017; Renna et al., 2011), inducing a decrease in DA levels, an increase in the strength of gap junction network and thus a shortening of the period of the retinal clock.

**Figure 7:**
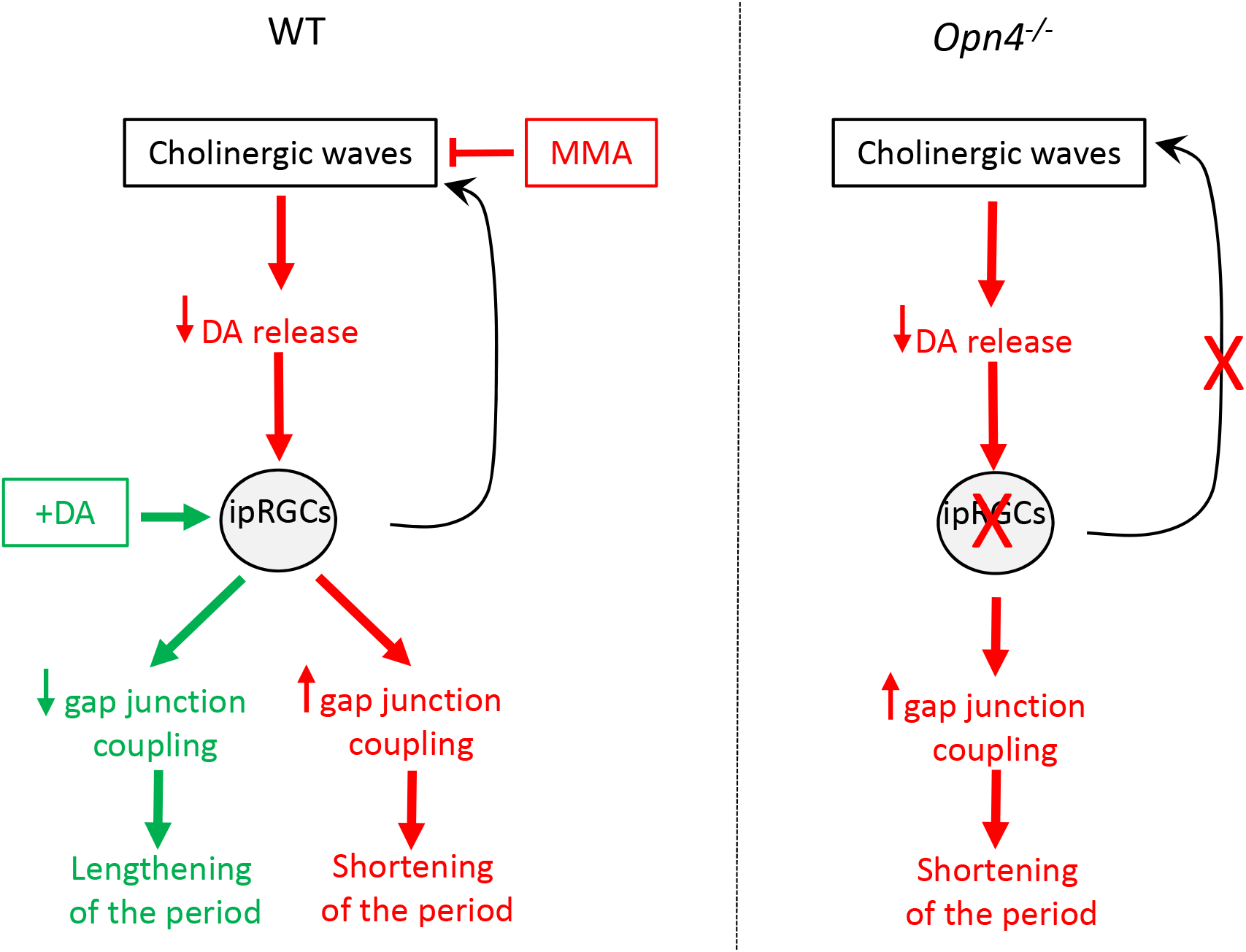
Model of the interaction between ipRGCs, cholinergic waves, and DA in early postnatal stages of retinal clock ontogenesis. In wild-type mice, DA supplementation decreases the extent of gap junction coupling among ipRGCs and other retinal cells, leading to a lengthening of the retinal clock period. When cholinergic waves are blocked, DA release is decreased, causing a robust electrical coupling between cells and a shortening of the period. The contribution of ipRGCs to retinal wave generation and dynamic (black arrow) is altered in the *Opn4*^*-/-*^ (right panel), leading to a decrease in DA levels, an increase in the strength of gap junction network, and thus a shortening of the period of the clock. ipRGC, intrinsically photosensitive retinal ganglion cell; MMA, mecamylamine acid.

The development of the retinal network is a gradual process beginning at postnatal ages, with thefirst development of clock capacity and later development of network properties. Elucidating the precise mechanisms that mediate the interplay between retinal waves, ipRGCs, DA, and retinal clock networks require future studies that will provide promising avenues for future research.

## Materials and methods

### Animals

Male and female *Per2*^*Luc*^ (Yoo et al., 2004) and *Opn4*^*-/-*^*::Per2*^*Luc*^ mice were housed in a temperature-controlled room (23 ± 1°C), under 12 h light/12 h dark cycle (12L/12D, light intensity around 200-300 lux) with food and water *ad libitum*. All animal procedures were in strict accordance with current national and international regulations on animal care, housing, breeding, and experimentation and approved by the regional ethics committee CELYNE (C2EA42-13-02-0402-005). All efforts were made to minimize suffering. Animals were used at embryonic day 18 (E18), postnatal days 1, 5, 8, 11, 15, 30, and in 2 month-old adults. The day of birth corresponded to P1.

### Retinal explant culture

For the embryonic stage 18, *Per2*^*Luc*^ pregnant mice were killed by cervical dislocation, and the embryos were isolated. Animals were sacrificed by decapitation between P1 and P8 and by cervical dislocation after P11, one hour before light offset at zeitgeber time 11 (ZT11). Eyes were enucleated and placed in Hank’s balanced salt medium (HBSS; Invitrogen) on ice. Retinas were gently isolated from the rest of the eyecup and flattened, ganglion cell layer up, on a semi-permeable (Millicell) membrane in a 35 mm culture dishes (Nunclon) containing 1.2 mL Neurobasal-A (Life Technologies) with 2% B27 (Gibco), 2 mM L-Glutamine (Life Technologies) and 25 U/mL antibiotics (Penicillin/Streptomycin, Sigma), incubated at 37°C in 5% CO2 for 24 h. From this step onwards, all manipulations of explants were performed under dim red light. After 24 h, at the projected ZT12, retinas were transferred to 1.2 ml of 199 medium (Sigma), supplemented by 4 mM sodium bicarbonate (Sigma), 20 mM D-glucose (Sigma), 2% B27, 0.7 mM L-Glutamine, 25 U/mL antibiotics (Penicillin / Streptomycin, Sigma) and 0.1 mM Luciferin (Perkin). Culture dishes were sealed and then placed in a Lumicycle (Actimetrics, Wilmette, IL, USA) to record the global emitted bioluminescence. All medium changes were performed under dim red light.

### Pharmacological treatments

For such experiments, 199 medium was supplemented with different molecules that remained in the medium throughout the entire recording without further medium changes: DA (50 µM), an inhibitor of DA synthesis alpha methyl-L-tyrosine (L-AMPT, 100 µM), an inhibitor of vesicular DA uptake reserpine (Res, 10 µM), a non-selective DA agonist apomorphine (Apo, 50 µM), a DA D1R antagonist SCH39166 (50 µM), a DA D2R antagonist L741626 (25 µM), a non-selective gap junction blocker CBX (100 µM) and mecamylamine, an acetylcholine receptor antagonist (MMA, 100 µM).

### Quantitative RT-PCR

Two retinas of wild-type and *Opn4*^*-/-*^ mice at P8 and P30 were dissected at ZT11, pooled, and stored at −80°C until RNA extraction and quantification. Total RNAs were extracted using Trizol reagent (Invitrogen) and reverse transcribed using random primers and MMLV Reverse Transcriptase (Invitrogen). Real-time RT-PCR was then performed on a LightCycler™ system (Roche Diagnostics) using the light Cycler-DNA Master SYBR Green I mix. Hypoxanthine ribosyl-transferase (*Hprt*) was used for the internal standardization of target gene expression. The efficiency and the specificity of the amplification were controlled by generating standard curves and carrying out melting curves. Relative transcript levels of each gene were calculated using the second derivative maximum values from the linear regression of cycle number versus log concentration of the amplified gene. Primer sequences were: *Hprt* sens ATCAGTCAACGGGGGACATA and reverse AGAGGTCCTTTTCACCAGCA; *D1R* sens CAGCCTTCATCCTGATTAGCGTAGGCG and reverse CTTATGAGGGAGGATGAAATGGCG; *D2R* sens CAGTGAACAGGCGGAGAATG and reverse CAGGACTGTCAGGGTTGCTA; *D4R* sens CGTCTCTGTGACACGCTCATTA and reverse CACTGACCCTGCTGGTTGTA; *D5R* sens CATCCATCAAGAAGGAGACCAAGG and reverse CAGAAGGGAACCATACAGTTCAGG.

### Data analysis

The period, the phase, and the amplitude of PER2::Luc oscillations were determined using SigmaPlot 12.5 software by fitting a linearly detrended sinusoidal curve oscillating around a polynomial baseline to the first three complete oscillations from each sample as previously described (Calligaro et al., 2019, 2020; Jaeger et al., 2015). The equation used was:

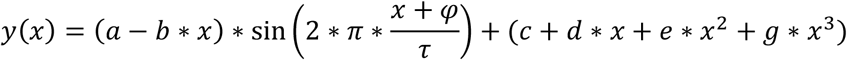

Where a is the amplitude, τ the period, and φ the phase. Rhythm onset was defined when the first complete bioluminescence oscillation was observed (first through and peak clearly distinguishable). The circadian time of the peak of the first oscillation, which refers to the phase in the current manuscript, was calculated based on a time reference (CT12), which corresponds to the time of medium change corrected by the endogenous period (Calligaro et al., 2019, 2020).

### Statistics

One-way ANOVA (Kruskal Wallis) is performed, followed by the *post-hoc* Mann-Whitney test (Statistica Software) in order to compare the period, the phase, and the amplitude of PER2::Luc signals, pharmacological treatments, and the relative expression of DARs during development. Two-way ANOVA, followed by the post-hoc Mann-Whitney test, was used to compare the period, the phase, and the amplitude between *Per2*^*Luc*^ and *Opn4*^*-/-*^*::Per2*^*Luc*^ mice during development. Data are represented as mean ± SEM.

## Acknowledgements

CK, HC, AJ, and CC performed the experiments. CK, HC, AJ, and ODB analyzed the data. CK and ODB conceived the study and designed the experimental plan. CK, NH, MB, and ODB wrote the manuscript. All authors took part in the revision of the manuscript and approved the final version. The authors do not have any potential conflict of interest. This research was supported by Rhône-Alpes CMIRA, USIAS, ANR Light-Clock. The funders had no role in study design, data collection, and analysis, decision to publish, or preparation of the manuscript.

---

**Supplementary figure 1:**
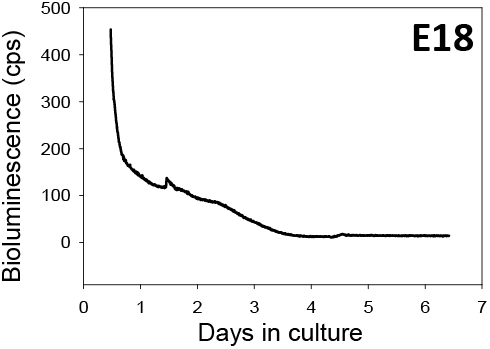
Representative bioluminescence recording of PER2::Luc retinal explants at embryonic day 18 (E18). No oscillations of PER2::Luc were detected in cultured retinal explants.

**Supplementary figure 2:**
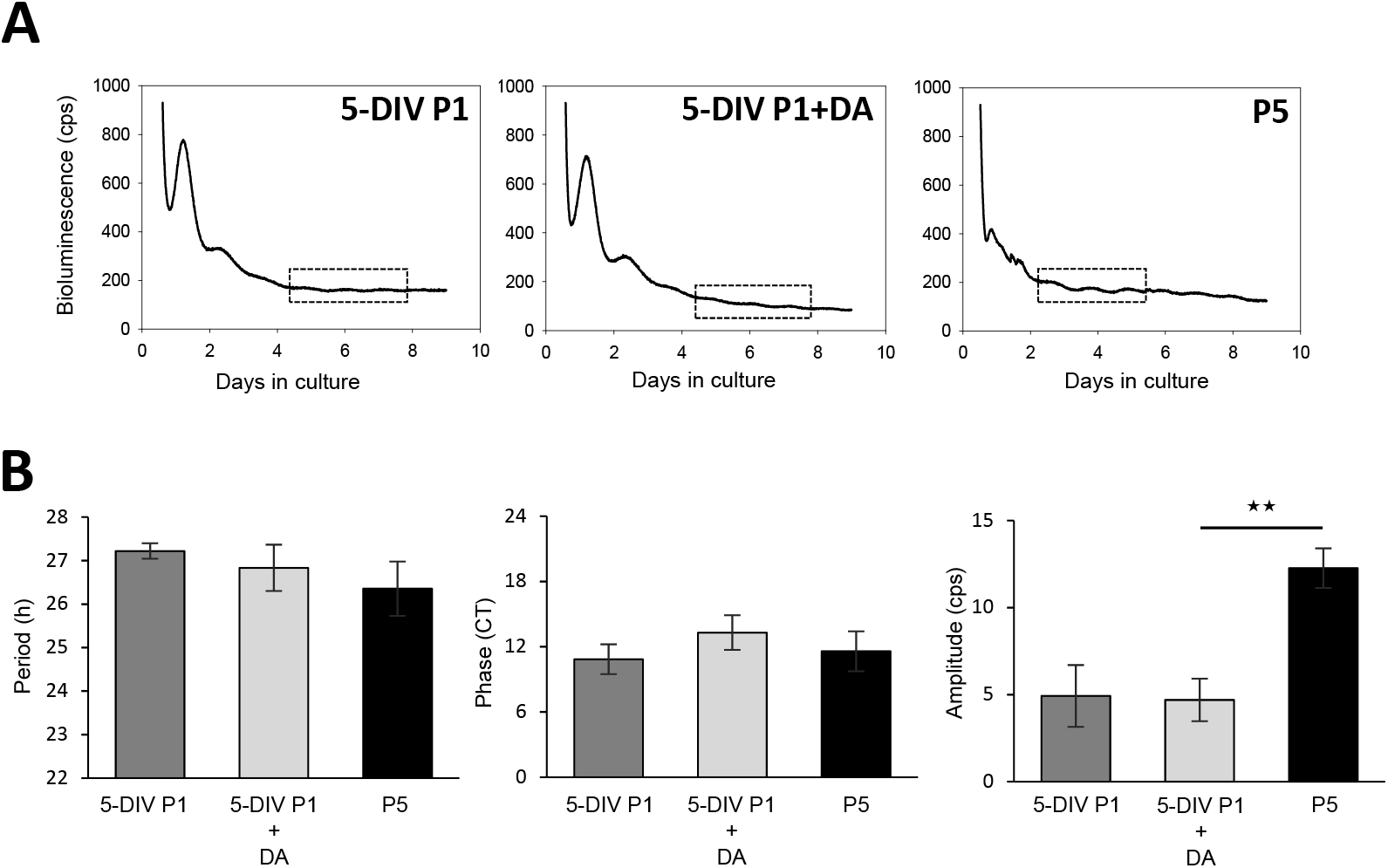
DA did not influence the onset of the spontaneous oscillations of PER2::Luc *in vitro*. **A-** Representative bioluminescence recording of PER2::Luc 5 days *in vitro* P1 cultured explants supplemented with DA (5-DIV P1 + DA). The dotted black rectangles correspond to the analysis windows, including the 3 first complete oscillations. **B-** Comparison of the means of the endogenous period, the phase, and the amplitude of 5-DIV P1, 5-DIV + DA, and P5 retinal explants. The total numbers of retinas used were: 5-DIV P1, n=5; 5-DIV + DA, n=4, and P5, n=6. Bars represent mean ± SEM (ANOVA, ** = p <0.01).

